# DrugForm-DTA: Towards real-world drug-target binding Affinity Model

**DOI:** 10.1101/2025.05.22.655461

**Authors:** Ivan Khokhlov, Anna Tashchilova, Nikolai Bugaev-Makarovskiy, Olga Glushkova, Vladimir Yudin, Anton Keskinov, Sergey Yudin, Dmitry Svetlichnyy, Veronika Skvortsova

**Author notes:** These authors contributed equally to this work (Ivan Khokhlov, Anna Tashchilova).

## Abstract

Drug-target affinity (DTA) prediction is a fundamental problem in drug discovery. Computational methods for predicting DTA can greatly assist drug design by decreasing the search space and reducing the number of protein-ligand complexes with low affinity. Currently DTA approaches often do not require protein three-dimensional (3D) structural information, which is often not accessible. In this study we present the DrugForm-DTA model, which uses only structure-less representations of ligand and protein. It is a Transformer-based neural network with protein encoding based on ESM, and small molecule ligand encoding obtained with Chemformer.

We evaluated the model on standard benchmarks Davis and KIBA, and revealed superior performance of DrugForm-DTA with best result for KIBA (MSE=0.117). Moreover, we developed a ready-to-use model using BindingDB dataset that was subjected to high-quality filtering and transformation. Overall, our method predicts drug-target affinity values with a confidence level comparable to a single in-vitro experiment. Also, we compared DrugForm-DTA against molecular modeling methods and revealed higher efficacy of the developed model for drug-target affinity predictions.

Our investigation provides a high accuracy neural network model with performance comparable to experi-mental measurements, filtered and reassessed BindingDB dataset for further usage, and demonstrates outstanding applicability of the proposed method for DTA prediction.

**Author summary:** Predicting drug-target binding affinity is a crucial task in computational drug discovery. Here we present DrugForm-DTA, a ready-to-use Transformer-based model which requires only protein amino acid sequence and ligand SMILES as inputs. The model shows excellent results at commonly used benchmarks.

We believe that the success in solving machine learning tasks depends on data quality and quantity, so the main focus of our work is the training data, but not a sophisticated neural architecture. We used BindingDB database as the source of experimental affinity measurements and prepared a refined and purified dataset, suitable for training machine learning models. Trained on this dataset, the DrugForm-DTA model demonstrates accuracy, comparable to a single *in-vitro* experiment. We also reveal that our model outperforms molecular modeling methods in estimating binding affinity. Both the prepared dataset and the trained model is published and freely available.

## Introduction

Developing a new drug that gains marketing approval is estimated to cost USD 2.6 billion, and the approval rate for drugs entering clinical development is less than 12% [1]. Classical laboratory screening technologies that measure the affinity of a small molecule to a target protein remain labor-intensive and expensive. Modern advances in computational methods and technologies, including the application of machine learning (ML) to chemical and biological research, make it possible to simultaneously expand the search space and to decrease the number of leader molecules.

One of the important tasks in drug design is predicting binding affinity for protein-ligand complexes (DTA, Drug Target Affinity). Modern approaches to computational prediction of DTA, related to the category of QSAR (Quantitative Structure–Activity Relationship), are based on various machine learning methods that have been actively developed in recent years, mainly neural network based. To train a DTA model, it is necessary to have reliable datasets with experimentally measured affinity constants (e.g., Ki, IC50). There are datasets traditionally used as a benchmark: Davis [2] and KIBA [3], but they contain information only on tens of thousands of protein-ligand complexes, which is insufficient for training an effective model. At the same time, large affinity measurement databases containing about a million records, such as BindingDB [4], PDBbind [5], BindingMOAD [6], are not directly ready for DTA training and need preliminary preparation and filtering.

Predicting drug-target affinity requires machine-readable representation of both protein and ligand data, and the choice of numerically representation of the protein and small molecule structures largely determines the success of training a DTA model. Small molecule structure description can be presented as a SMILES [7], a molecular graph, one of the pre-calculated descriptors (ECFP [8], Morgan [9], etc.), or even a 2D-image of the molecule [10, 11]. Pre-calculated descriptors convert a molecule structure into a numerical vector of fixed dimension, which entails informational loss, so they are rarely used as a full-volume representation of the molecule. However, some DTA models, like FingerDTA, utilize these descriptors [12]. Besides, the graph form is the most natural representation of a molecule structure. This approach was used in the development of the following models: DGraphDTA [13], MGraphDTA [14], DoubleSG-DTA [15], GraphDTA [16], HSGCL-DTA [17]. However, using graph representation leads to a number of complications in neural network architecture. The SMILES notation is ultimately the same molecular graph, expanded as a string and allows applying the neural network approaches from the NLP field (Natural Language Processing), mainly the transformer-like neural networks. As a result, the SMILES representation of a small molecule is increasingly used in the field (DeepDTA [18], ProSmith [19], MRBDTA [20], MFR-DTA [21]). Some studies combine several representation methods in one multimodal neural network to gain some advantage in the representation completeness (MultiScaleDTA [22], HGRL-DTA [23], 3DProt-DTA [24], BiComp-DTA [25], MSF-DTA [26], HGTDP-DTA [27]).

Protein encoding methods also vary. For example, the FingerDTA [12] model uses a fairly simple approach -the Word2Vec model [28]. The HGRL-DTA [23], 3DProtDTA [24], MultiScaleDTA [22] models use a 3D-structure generated by AlphaFold [29]. In the ProSmith [19] work authors used a primary amino acid sequence as text. To date, some of the most notable DTA models are ProSmith [19],

DeepDTA [18], and HGTDP-DTA [27]. The ProSmith DTA model [19] uses a multimodal Transformer Network to process an amino acid sequence together with a SMILES string as an input pair. This approach estimates conjunction between the protein and ligand structures attempting to predict their structural and therefore functional interaction. The ESM-1b [30] model is used to represent proteins, and the ChemBERTa2 [31][31] model to represent ligands. The ProSmith model was pre-trained for six epochs on the BindingDB dataset [4], which contains affinity values for about a million protein-ligand pairs. Then the obtained neural network parameters were used to continue training the model on the Davis benchmark dataset [2]. The authors used this approach due to the insufficient amount of data in the Davis dataset for training (about 30,000 records). As a result, at the time of publication, the ProSmith model performed the best at the Davis benchmark. Another approach represent a protein and a small molecule is used in the DeepDTA model [18] and based on the integer encoding for both the protein and the ligand (e.g., C:1, H:2, N:3, etc.) in a convolutional neural network (CNN) with two separate inputs. The HGTDP-DTA affinity model [27] processes a ligand, represented as SMILES, and a protein as the amino acid sequence were transformed into molecular graphs to obtain numerical vector representations (embeddings) with a GCN. Then, both molecular graphs were combined into a single feature space using GCN and Transformer.

In this work we aimed to create a new efficient and accurate deep learning model for predicting drug-target affinity (Fig 1). We developed a new approach, called DrugForm-DTA, which uses only the primary amino acid sequence and the SMILES as inputs. Our model uses a fairly simple neural network architecture based on Transformer, shifting the focus from the complexity of the neural network structure to the quality of the model training procedure and the quality of the training dataset. The DrugForm-DTA model was tested on standard benchmarks - Davis and KIBA, and compared with other top-tier DTA methods for the year 2024: MultiscaleDTA [22], HGRL-DTA [23], MFR-DTA [21], DGraphDTA [13], BiComp-DTA [25], MSF-DTA [26], GraphDTA [16], DoubleSG-DTA [15], 3DProtDTA [24], HGTDP-DTA [27], MGraphDTA [14], HSGCL-DTA [17], MRBDTA [20], FingerDTA [12]. In order to obtain a highly accurate DTA model, applicable for real-world usage, we prepared a large and thoroughly processed training dataset based on the BindingDB database. BindingDB is one of the largest public databases which contains more than 2.8 million experimental measurements of affinity constants. We provide the prepared training dataset along with its test part, separated with a combination of cold target and drug scaffold splits. We trained a DTA model on the prepared BindingDB dataset and proved high efficiency in predicting the affinity constants of a small molecule to a protein. The DrugForm-DTA model demonstrated higher applicability compared to approaches that require the presence of the 3D protein structure, such as molecular docking, molecular mechanics, and semiempirical methods of quantum chemistry.

**Fig 1.**
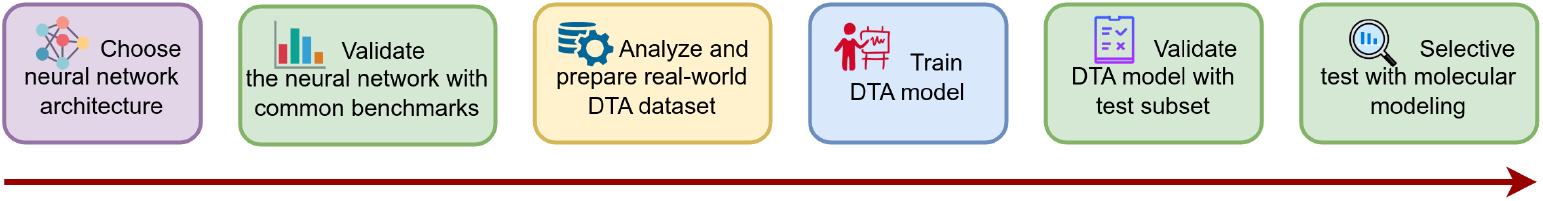
Short pipeline of the work.

## Results

### DTA neural network architecture

Neural networks are one of the most advanced approaches to solving QSAR problems. Transformer-like neural networks [32] effectively work with string sequences of different lengths. This is very convenient in the DTA problem, since the ligand and protein molecules are represented by a SMILES and an amino acid sequence, respectively. Here we intensively used neural network architecture for solving the DTA problem taking into account recent achievements in the field [19, 30, 33]. We tested two neural network structures. The first one is a BERT-like network (encoder-only), where the input sequence is formed by concatenation of ligand and protein embeddings with a separator. This network is similar to ProSmith [19], but uses a different molecule encoder - Chemformer [33] (combined variant, Fig 2A), which seems more interesting to us compared to ChemBERTa2 [31] used there. Chemformer performs significantly better than ChemBERTa2 on the MolNet benchmark [34].

**Fig 2.**
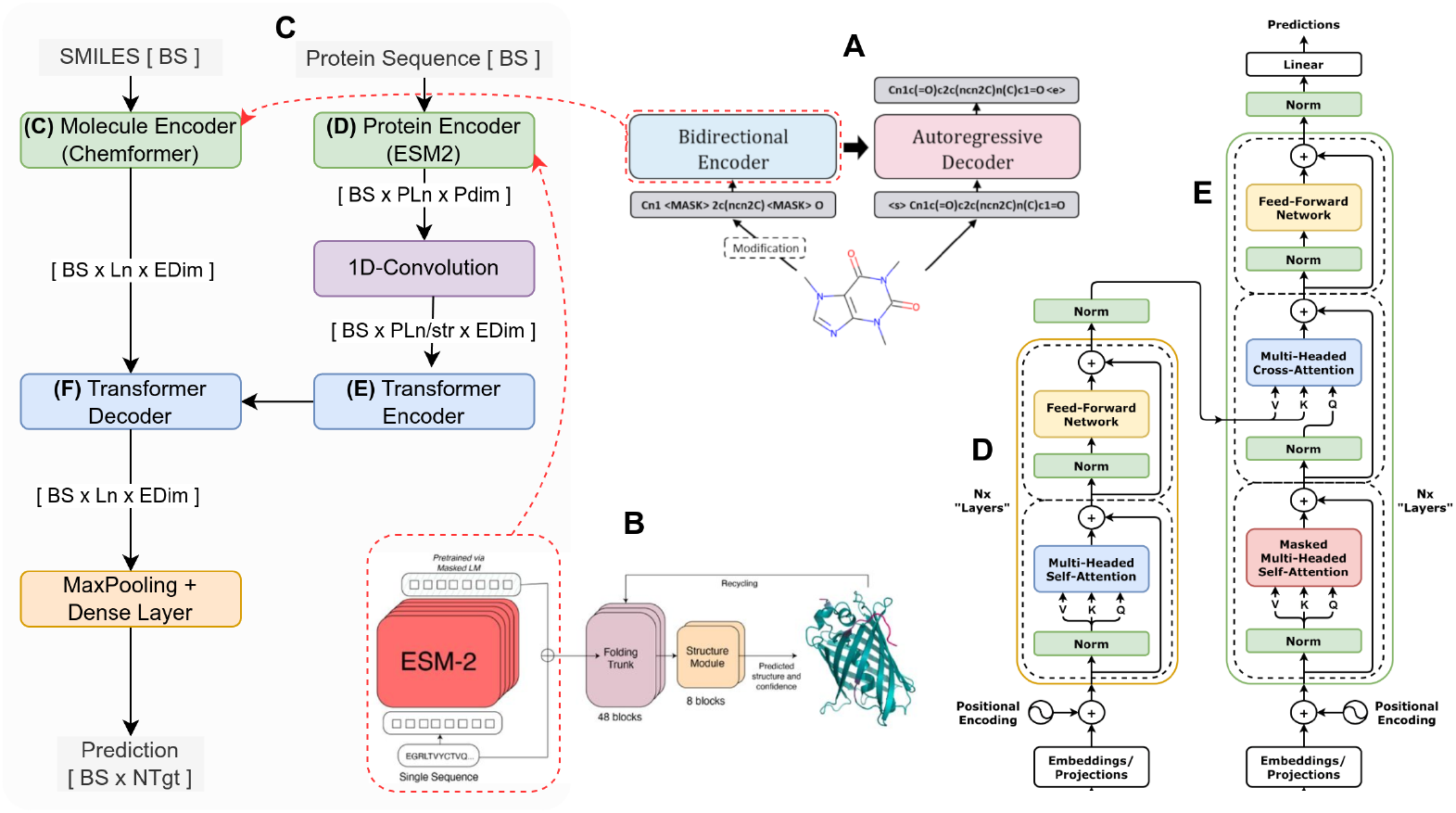
Proposed neural network for DTA prediction. A: Using the Chemformer model (combined variant) as a non-trainable molecule embedding generator. Adopted from [33]. B: Using ESM model (esm2 t30 150M UR50D variant) as non-trainable protein embedding generator. Adopted from [30]. C: DrugForm-DTA neural network architecture. The decoder block (F) takes the molecular embedding as input, while conditioning with protein embeddings through the encoder block (E). The 1D-convolution layer is fixing the difference between molecule and protein embedding dimensions. D, E: Vanilla transformer encoder and decoder. Adopted from [32].

The ESM model (esm2 t30 150M UR50D, Fig 2B) is used to generate protein embeddings [30], and the ligand and protein embeddings must have the same dimension for concatenation. The ligand embedding has a dimension of 512, and the protein embedding has a dimension of 640, so it must be converted to the same dimension. To this end, we applied a 1D-convolution that performs the same function instead of a linear layer like in the ProSmith work [19].

A minor disadvantage of our initial network structure is the increased size of the encoder input (protein & molecule) due to self-attention *O*(*N* ^2^) complexity. Proposed network structure (Fig 2C) is a vanilla transformer consisting of an encoder (Fig 2D) and a decoder (Fig 2E). Protein embedding is fed to the encoder and determines the decoder behaviour by encoder-decoder attention. The final network requires fewer calculations than the initial one, since 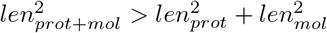. We have performed experimental measurements, and for SMILES with a length of 70 symbols and a protein of 500 amino acids, the initial model spent 34 seconds, and the final model just 32 seconds, which is 6% less. For a larger protein with a length of 1500 amino acids, the calculation time for the initial model was 140 seconds, and for the final one 131 seconds, which is 7.5 percent less. We conducted all tests with 1000 iterations and a batch size of 16.

Thus, the neural network accepts a SMILES string and a primary amino acid sequence as inputs. The SMILES string is processed with the Chemformer encoder, and the protein sequence is processed by the ESM encoder. Transformer decoder block contains 6 layers, and Transformer encoder block contains only one layer, because features are already extracted by ESM. Max Pooling layer in combination with Dense layer converts decoder embeddings into a vector of output values, representing affinity between a given ligand and protein.

### DTA benchmarks

The proposed neural network architecture was tested on the traditionally used Davis [2] and KIBA [3] benchmarks. The KIBA and Davis datasets require a regression approach for the evaluation of models, so the main benchmarking metric is MSE (Mean Squared Error). We used RMSE (Root Mean Squared Error) metric, because it numerically corresponds to the range of target values. To compare DrugForm-DTA with other models, the values were recalculated into MSE. Besides MSE we also used the C-index to compare performances across models.

First, we estimated correspondence between the predictions of the trained model and target values on the test subsets. Both for KIBA (Fig 3A) and Davis (Fig 3B) benchmarks our results indicate a moderate correlation between the model predictions and the target values. Moreover, the points clustered on the left side of the plot (Fig 3B) correspond to the records with the target value pKi=5, which dominate in the Davis dataset. So, this indicates a problem with benchmarking datasets that require further investigations in order to properly interpret predictions and evaluate models.

**Fig 3.**
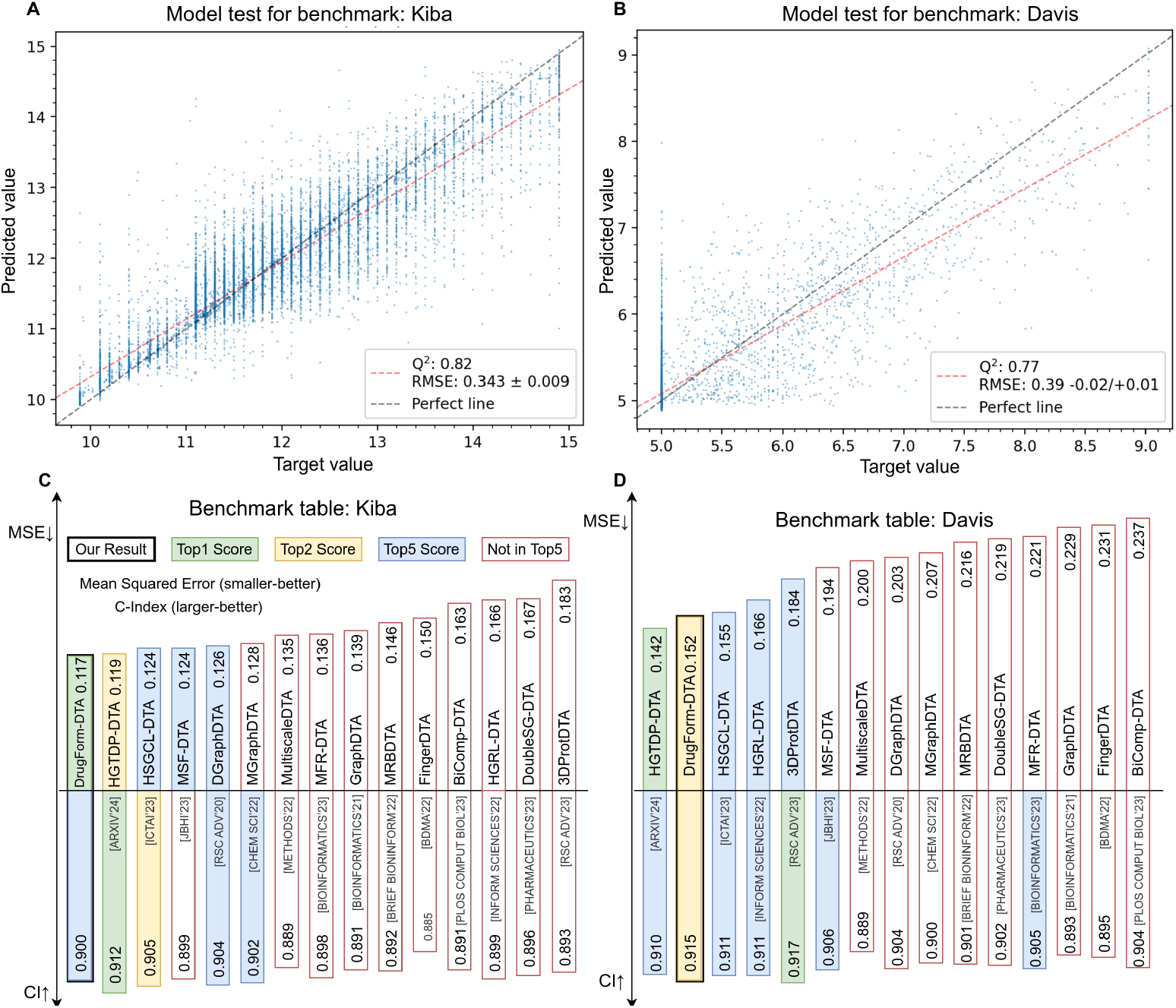
Performance of the DrugForm-DTA model on the Davis and KIBA datasets. A: Visualization of the prediction on the KIBA test subset. There is a moderate correlation between the model predictions and the target values. The dataset is dominated by rounded values. B: Visualization of the prediction on the Davis test subset. C: Comparison of prediction performance across DTA models on KIBA benchmark. DrugForm-DTA ranked first in MSE metric with a value of 0.117. D: Comparison of prediction performance across DTA models on Davis benchmark. DrugForm-DTA ranked second with MSE=0.152.

Next, we compared DrugForm-DTA model performance with other published DTA models: MultiscaleDTA [22], HGRL-DTA [23], MFR-DTA [21], DGraphDTA [13], BiComp-DTA [25], MSFDTA [26], GraphDTA [16], DoubleSG-DTA [15], 3DProtDTA [24], HGTDP-DTA [27], MGraphDTA [14], HSGCL-DTA [17], MRBDTA [20], FingerDTA [12]. Despite previously mentioned problems in the Davis dataset, we benchmarked DrugForm-DTA with other methods using the same data to evaluate model performance. The DrugForm-DTA model achieved the top 1 performance on the KIBA benchmark (Fig 3C) (MSE=0.117, C-index=0.912), and for the Davis benchmark (MSE=0.152, C-index=0.915) our model is the second best (Fig 3D). We also evaluated the C-index metric, and DrugForm-DTA outperforms HGTDP-DTA. However, the 3DProt-DTA approach has the best C-index metric across all models but significantly inferior in MSE (MSE=0.184).

Overall, we benchmarked the DrugForm-DTA model against other approaches,and revealed high performance of our method. We used both MSE and C-index metrics that can be discordant in ranking across methods. Also, we point to the problems in the benchmarking datasets that require further investigations in order to properly interpret model quality.

### BindingDB preparation

Building an efficient and accurate affinity model requires a high-quality, consistent dataset. So, we performed additional steps to prepare the datasets and train model relying on the most accurate instances. We obtained binding affinity values from the public BindingDB database [4] to train DrugForm-DTA, which contains more than 2.8 million experimental measurements (1,238,443 ligands and 6,523 proteins). However, we discovered several issues deteriorating quality of the training data: existence of threshold values, multiple measurements for the same protein-ligand complexes, lacking full information about proteins. We defined an approach to solve the issues.

We noticed that many experimentally measured binding affinity values are reported in BindingDB (Fig 4A) as a threshold value (for example, *>*5, *>*6). We further assume a serial dilution method behind this issue. Excluding these records leads to information loss, and simply removing the thresholds causes significant deviations in accuracy (about half order of magnitude). As a compromise we replace threshold values with a hypothesis of a real value by modifying reported threshold affinity value by 1/2 order away from the threshold Moreover, this way we also achieve smoothing of the measurements reported as threshold values and anomalously overrepresented. Thus, modifying records with the same value prevents the neural network from learning this value instead of approximating the hypersurface of the function being optimized [35].

**Fig 4.**
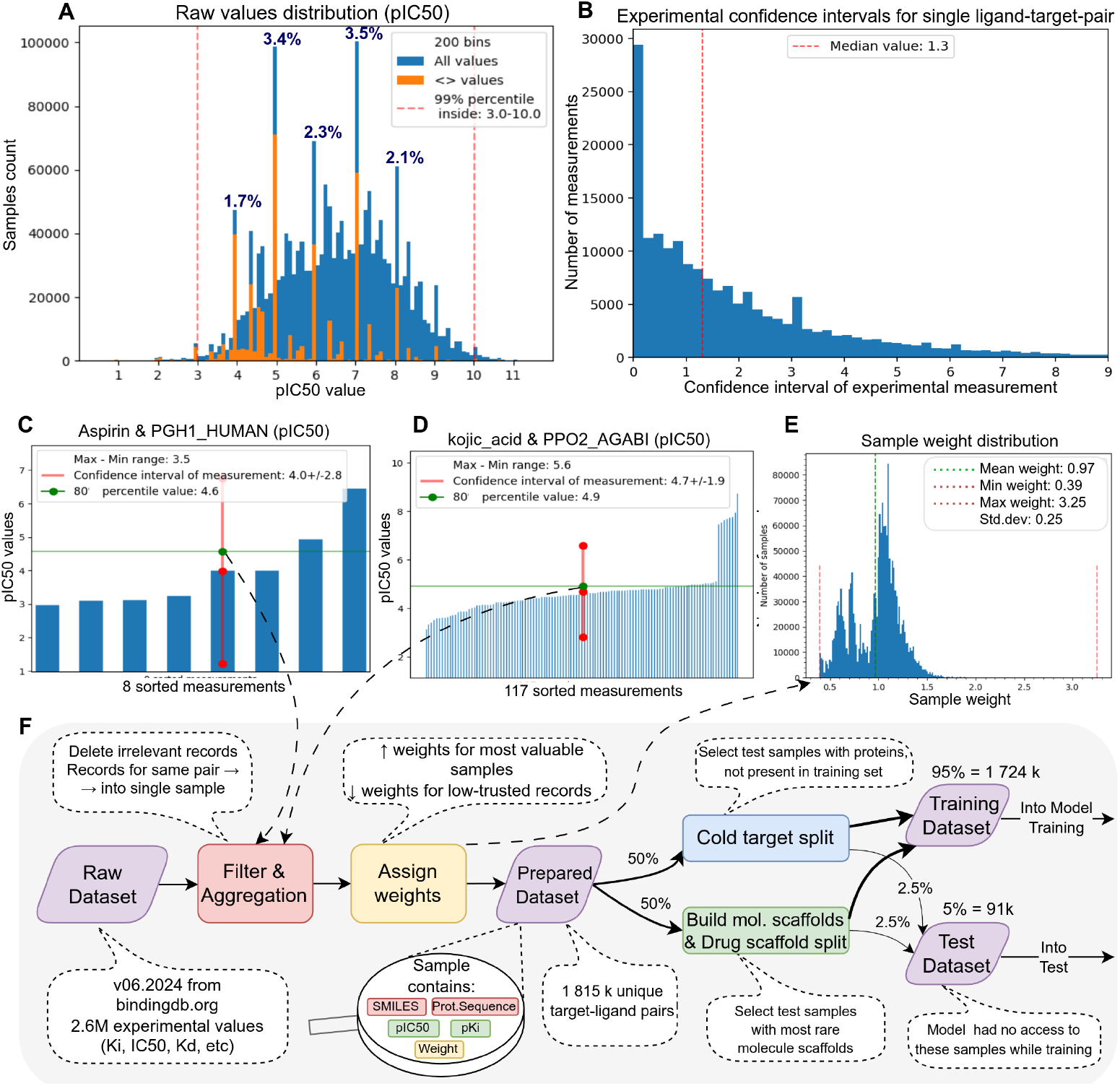
Preparation of the BindingDB dataset. A: Distribution of raw values in the dataset (using pIC50 as an example). Orange color shows how many values are specified with the ¡ or ¿ threshold. Records with round values dominate. Their occurrences in percentages of the whole dataset are indicated above the peaks. B: Histogram of the confidence interval values of the experimental measurement of affinity constants for individual protein-ligand pairs. Zero values are for pairs with identical values. Even taking them into account, the median value is 1.3 orders of magnitude. C: An example of aggregating several measurements for one protein-ligand pair: the 80th percentile value (4.6) represents a soft maximum that is robust to outliers and will be used in training. Experimental confidence interval is 4.0+/-2.8. D: An example of robustness to outliers: a group of measurements with results significantly exceeding the norm is discarded. 80th percentile value (4.9) is close to average value (4.7) and quite far from maximal value (8.7) E: Distribution of example weights in the final dataset. Large weights were assigned to the most significant or reliable records, small weights to the least reliable ones. The example weight is taken into account during the training process. F: Schematic diagram of the original BindingDB database transformation into training and test datasets. The full procedure starting with downloading raw data from the BindingDB website to forming the training and test datasets is shown.

One of the important issues is the presence in BindingDB of several different measurements for 43% of the unique protein-ligand complex. For each protein-ligand pair with two or more measurements we estimated the confidence interval (95%) (Fig 4B) and difference between the maximum and minimum values (Fig B in S1 Text). Estimating average confidence interval as 1.3 order and average min-max difference as 0.9 order looks embarrasing and demonstrates inconsistency of experimental data. For example, available records in BindingDB for such a basic complex as Aspirin & PGH1 HUMAN revealed values in quite a wide range from 3 to 6.5 (Fig 4C). Existing approaches [36] to deal with multiple measurements suggest taking of mean or maximum value. The motivation for taking the maximum value instead of mean value is that an experimental error is more likely to decrease the affinity than to increase it [37] (see the Binding Affinity Experimental Measurement Issues chapter in S1 Text), but taking strictly maximal value is not robust to outliers. As a compromise we use the 80 percentile value, which behaves like a soft maximum but robust to outliers.

An example of such robustness to outliers is the aggregation of pIC50 values for the pair kojic acid & PPO2 AGABI (Fig 4D), where the 80-percentile is close to the average and ignores the group of high values that stands out from the general mass. We also note that in practice we take a weighted percentile. The weights (importance) of records are formed from various factors which make an individual record more or less valuable (Fig 4E).

We noticed that the BindingDB dataset sometimes provides only the binding site part instead of the entire protein sequence, and sometimes the sequence contains incorrect symbols that do not exist in the amino acid dictionary. Therefore, the amino acid sequences were obtained from the UniProt database directly [38]. In addition, information of the protein host organism was extracted from UniProt.

The overall BindingDB filtration and transformation procedure (Fig 4F) allowed to preserve as much data as possible without significant contamination of the dataset. The final dataset contains 1,739,873 protein-ligand records (5,251 proteins, 1,020,614 ligands) with parameters: amino acid sequence, SMILES, pKi, pIC50, weight.

The final step of preparing a dataset to model training is splitting it into training and test parts. We assigned 5% of the dataset as the test part, or more than 90 thousands records. It is comparable to the size of the full dataset in some works on the preparation of BindingDB (ELECTRA-DTA [39] 144 thousands protein-ligand pairs, BiComp-DTA [25] - 42 thousand, SAM-DTA [40] - 232 thousands), and is enough for creating a high-quality DTA model. Random split is considered suboptimal [34, 41, 42], so we formed one half of the test set with cold target split, and another half with drug scaffold split.

Overall, filtering and transformation of the BindingDB data represents part of our approach for training of the high performance binding affinity model. We also consider such data preparation as one of the most important steps towards a model able to accurately predict affinity of the ligand-target interactions. Further steps include computational techniques that we used in order to train and evaluate the DrugForm-DTA approach.

### Training and performance of the DrugForm-DTA model

After preparing the optimized BindingDB dataset, we trained the DrugForm-DTA model (Fig 5A) using a multitask approach to produce two parameters at once: pKi and pIC50. Also, we used the test part of the BindingDB (90,763 records, see BindingDB Preparation) for independent model evaluation. DrugForm-DTA model uses SMILES and primary amino acid sequence as input, and each string passes through corresponding encoder (Chemformer or ESM) to obtain necessary embeddings (Fig 5B). One of the key elements in the training of the full DrugForm-DTA model is cross-validation technique and averaging of the submodel prediction results (Fig 5C). The test part of the dataset was isolated and did not participate in training. The training dataset was also divided into training and validation subsets in four parts without crossing the validation sets (classical K-fold CV [43]). Each submodel learned (see Materials and Methods) on the 3/4 of the full training set, and validated at 1/4 of the full training set. The validation subset is used as overfitting control and model selection criterion. This approach improves the predictive power of the model at the cost of additional computations. The neural networks used in the model are computationally heavy, but the embeddings produced by the non-trainable ligand and protein encoders do not change over time and were cached. Thus, the additional cost of introducing four submodels is less than four times. We trained a full DrugForm-DTA model using four NVidia A100 GPU for 30 days.

**Fig 5.**
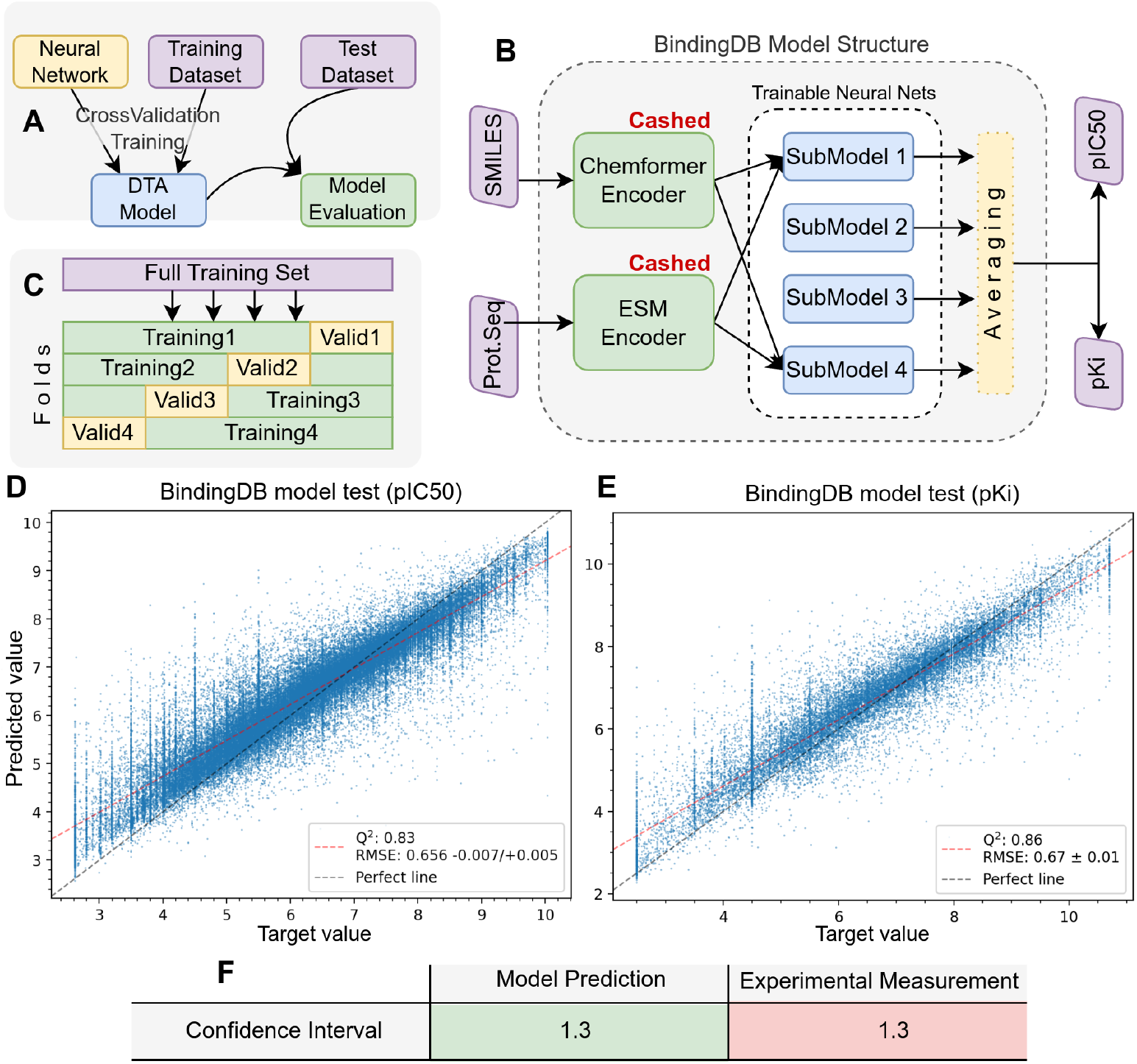
DTA model trained on the BindingDB dataset. A: Splitting the original dataset into training and test subsets. B: Structure of the DTA model. The model consists of several submodels, one for each cross-validation fold. The results of the submodels are averaged to obtain a consolidated value. The protein and ligand encoders are cached and calculated once. The model takes the primary amino acid sequence and SMILES as input and returns two values at once: pKi and pIC50. C: Splitting the training dataset into a group of training and validation subsets by K-fold cross-validation: individual parts of the dataset alternately become training or validation. D, E: Evaluation of the quality of the trained model using the test set. RMSE(pKi)=0.66, RMSE(pIC50)=0.67. There is an observable correlation between the model predictions and the target values. F: Comparison of confidence intervals of experimental measurement and model calculation.

The trained model was applied to the test set. The results of applying the model to the test set are presented in scatter plots separately for pIC50 (Fig 5D) and pKi (Fig 5E). The accuracy of the predictions on the test set is RMSE(pIC50)=0.66 and RMSE(pKi)=0.67. These values are close to each other, so we can estimate the model accuracy based on the worst one: RMSE=0.67 (MSE=0.45). The average confidence interval of the model prediction on a large dataset can be estimated as *CI_calc_* = 1.96 *· RMSE* = 1.3 (Fig 5F, see Materials and Methods). We also calculated the confidence interval of the single experimental affinity measurement at data preparation step: *CI_exp_* = 1.3 (Fig 4B). Thus, accuracy of the model predictions is comparable with a single experimental measurement.

### DTA model selective test

#### Test samples selection

Next, we performed additional computational analysis using molecular docking, MM-GBSA calculations and semiempirical methods of quantum chemistry to estimate agreement of the experimental values with affinity predictions from the DrugForm-DTA and molecular modeling. To this end, we selected cases from the BindingDB test set not used for model training. All selected records cover target proteins with therapeutic effects, available multiple ligands with a wide range of affinity values, and X-ray diffraction protein structure exists in PDB [44].

Selected cases include targets for the insomnia treatment: melatonin receptor 1A (MTR1A), melatonin receptor 1B (MTR1B), orexin/hypocretin receptor type 1 (OX1R), orexin receptor type 2 (OX2R). The crystallographic structures of these target proteins have low resolution (2.6-2.8Å) and missing amino acid residues but far from the active site. Also, we selected antitumor activity ligands of the epidermal growth factor receptor (EGFR). High resolution (1-1.5Å) crystallographic structure of the EGFR is currently available from the PDB database but contains a significant number of missing amino acid residues. Another example that we considered is blood coagulation factor XIa (FA11) that is used as a target for the development of anticoagulants. Although the structure has a high resolution (1.6Å), the amino acid sequence is represented only by the heavy chain and a small region of the light chain. The sodium- and chloride-dependent glycine transporter 1 (SC6A9) is selected as target for the schizophrenia treatment, the protein crystallographic structure has a low resolution (3.4Å) and about 90 amino acid residues missing at the N-terminus of the chain.

Overall, we selected 20 ligands for each protein with a range of affinity values from 2 to 10 excepting MTR1B with only 7 available ligands in the test set. We included this receptor to analyze the model behavior for proteins belonging to the same family. Our final evaluation dataset contains 127 protein-ligand pairs and used further to estimate agreement between DTA model predictions and binding energy obtained with molecular modeling methods.

#### Selective test metrics

Next, we applied molecular docking, molecular mechanics and quantum chemical calculations for each protein-ligand complex. We develop an algorithm to combine the results of ligand-protein binding measurements for the three approaches Fig 6A).

**Fig 6.**
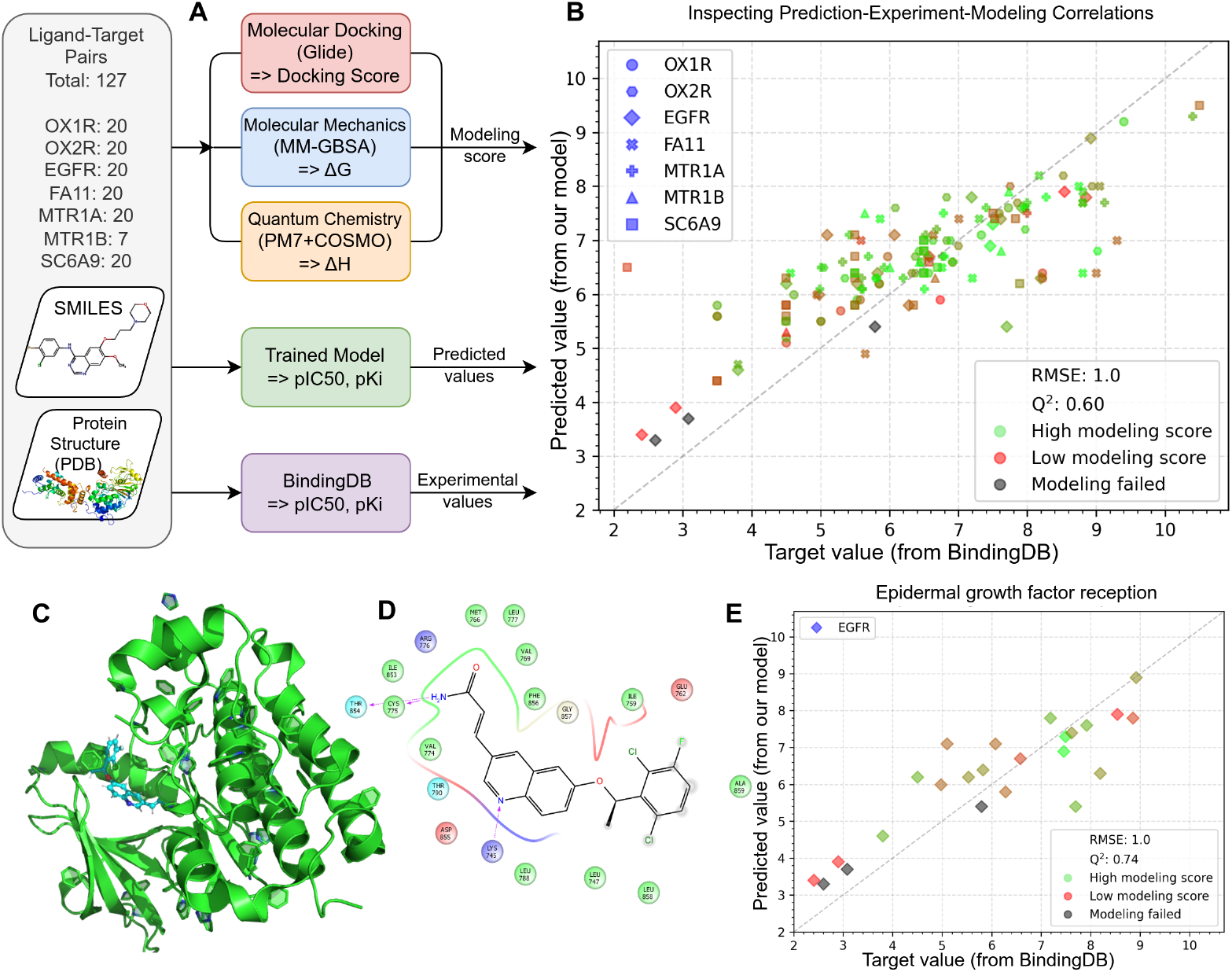
Molecular modeling. A: Flow chart of the detailed testing of single protein-ligand pairs. The modeling score was calculated as the average of the ranked protein-ligand interaction energies obtained by molecular docking (Glide), molecular mechanics (MM-GBSA), and quantum chemical calculations (PM7+COSMO). B: Scatter plot of the affinities predicted by the DTA model, experimentally measured affinities, and predicted by molecular modeling for 127 protein-ligand complexes. The X-axis shows the experimental affinities, the Y-axis shows the DrugForm-DTA predicted affinities, and the color shows the calculated modeling scores. C: Scatter plot of affinities predicted by DTA model, experimentally measured and predicted by molecular modeling methods for EGFR and 20 ligands. Experimental affinities are shown on the X-axis, DrugForm-DTA predicted values are shown on the Y-axis, and calculated modeling scores are shown in color. All protein-ligand pairs with modeling failed (marked in gray) refer to the EGFR protein. D: 3D structure of EGFR in complex with the ligand, the ligand position is obtained as a result of molecular docking. E: Position of the ligand in the active site of EGFR. Hydrogen bonds formed with amino acids of the protein are shown with pink dashed arrows.

We performed molecular docking for 127 protein-ligand pairs using the Glide software [45]. Atomistic protein models were suitably prepared based on the crystal structures from the Protein Data Bank [44]. Three-dimensional structures of ligands were constructed taking into account the isomerism and charge states of the molecule, and to approximate the protein-ligand interaction energy we used Glide docking score. Also, for all protein-ligand pairs we applied MM-GBSA analysis (Prime) to calculate ligand-binding (Δ*G_bind_*) affinities [46] using ligand position in the protein active site from the Glide docking. In order to obtain the most accurate positioning of the ligand, we applied quantum chemistry methods (PM7+COSMO) [47, 48] that optimized ligand positioning in the protein active site and allowed a more accurate estimation of the binding enthalpies. Finally, we introduced a modeling score that represents the mean of the MM-GBSA, Glide and quantum chemistry predicted binding (see Materials and Methods). Obtained modeling scores are continuous, but we split them into classes with low, medium and high modeling scores, and a category where one or all molecular modeling approaches did not yield a result.

#### Selective test results

We have compared affinities from BindingDB with DTA model predictions, and molecular modeling score (Fig 6B). We report a low correlation between the modeling score and the experimental affinity values with Spearman correlation value *C_Mod_*=0.20, while DTA model predictions have significantly higher Spearman correlation value *C_DT A_*=0.76. In particular, protein-ligand complexes with low modeling scores often have high binding affinity according to the experimental measurements (see Data Availability section).

For the EGFR case (Fig 6C) we revealed a cluster of high modeling score values in the high affinity region, but there are also low modeling scores (*C_Mod_*(EGFR)=0.18, *C_DT A_*(EGFR)=0.80). As an example, the positioning of a low-modeling score ligand in the EGFR active site was analyzed (Fig 6D). Glide docking results (Fig 6E) indicates the ligand forms three hydrogen bonds with the protein, but the docking score values and other modeling methods were low. All protein-ligand complexes with failed molecular modeling belong to the EGFR protein (Fig C in S1 Text) due to the protein fragment PDB structure (protein is partially crystallized and more than 70% of the amino acid sequence is missing). Our results exemplify the limitations of the molecular modeling methods for binding prediction due to the absence of a complete crystal structure of the target protein with good resolution.

We performed a detailed comparison of the molecular modeling results with the experimental and predicted affinity for each of the tested proteins. In particular, modeling of MTR1A (Fig D in S1 Text) and OX2R (Fig E in S1 Text) binding revealed high modeling score values for almost all ligands from the test set, even for those with lower experimental values, that leads to low correlation values (*C_Mod_*(MTR1A)=-0.03, *C_DT A_*(MTR1A)=0.80, *C_Mod_*(OX2R)=-0.26, *C_DT A_*(OX2R)=0.70). Thus, molecular scores are poorly suitable to distinguish between high and medium affinities (pKi/pIC50 in range from 5 to 8). Modeling of ligands binding with OX1R (Fig F in S1 Text, *C_Mod_*(OX1R)=-0.04, *C_DT A_*(OX1R)=0.84) also shows no correlation with experiment and reveals the need for not only a sufficiently complete, i.e. containing all amino acid residues, crystal structure but also need for a good resolution (*<*1.5Å). Both receptors have missing amino acid residues only at the end of the chain, far from the active site, but have a low resolution (*≈*2.8Å). For FA11 (Fig G in S1 Text, *C_Mod_*(FA11)=0.06, *C_DT A_*(FA11)=0.55) we have high-resolution crystal protein structure (*≈*1.6Å), but the modeling scores do not fully correlate with experiment, that is supposedly caused by missing a significant part of the amino acid residues of the light chain.

A large number of ligands with low modeling scores are also observed for protein–ligand complexes with SC6A9 (Fig H in S1 Text, *C_Mod_*(SC6A9)=0.27, (*C_DT A_*(SC6A9)=0.70), which is one of the best modeling results along with MTR1B (Fig I in S1 Text, *C_Mod_*(MTR1B)=0.40, (*C_DT A_*(MTR1B)=0.54).

Thus, DrugForm-DTA model predictions correlate with experimental values significantly better than modeling scores. Moreover, our model takes only the primary amino acid sequence, while molecular modeling methods require spatial structure of the protein. Molecular modeling is useful for positioning the ligand in the protein active site, but modeling scores do not display the affinity itself.

## Discussion

The main result of this work is the creation of DrugForm-DTA, a new trained model for predicting the affinity of interaction of a ligand molecule with a target protein, capable of predicting the values of pKi and pIC50 constants for any protein-ligand complex. The model allows predicting the affinity of a small molecule to a protein with a degree of confidence comparable to a single *in-vitro* experiment. The neural network architecture of DrugForm-DTA was preliminarily tested on standard benchmarks and showed suitability for full-fledged use. DrugForm-DTA uses a fairly simple neural network architecture based on Transformer, while the focus is shifted from the complexity of the neural network structure to the quality of the model training procedure and the quality of preparing datasets for training. We are convinced that it is the quality of the data, and not the complexity of the neural network architecture, that is crucial for a successful machine learning model. The used neural network architecture is quite simple and has potential for improvement. Following the ideas of the authors of other works mentioned above, it makes sense to try several input modalities for representing the ligand, for example, graph or descriptors. However, we believe that the main potential for improving the quality of the model depends on the high-quality training dataset. Indeed, the development of the model for predicting the affinity of protein-ligand complexes required additional preparation of datasets, since the use of available datasets was limited due to insufficient unification of data filling.

When evaluating the model performance, we revealed that choice of benchmarks was very limited and interpretation issues may arise. For the Davis benchmark dataset (Fig A in S1 Text), we found that almost 3/4 of the records in the dataset (71%) are 10*µ*M (pKi=5). The reason behind this issue is unclear and requires deeper investigation, but it limits the use of the dataset as a reference regression benchmark. In this regard, high quality benchmark datasets are necessary. We suggest that a high-quality benchmark dataset should be large and various enough to provide a good approximation of the model accuracy. A procedure similar to the introduction of example weights in the BindingDB-based training dataset considered in this paper can be used to select the most reliable examples. We suggest the reference data set should include only protein-ligand complexes where a large number of experimental measurements come from various teams and reported values are exact with low spread across measurements rather than a threshold. Our analysis follows that the KIBA dataset (117,657) is significantly larger compared to Davis (25,772), and the values are distributed in KIBA compared to Davis.

The high-quality and large dataset is a major factor influencing model performance and its applicability to real cases. The BindingDB database is one of the largest sources of experimental affinity constant values, and we used it to prepare our training dataset. It is a raw database, which contains threshold values, missing fields, partial amino acid sequences, anomalously overrepresented values, and also for half of its protein-ligand pairs it provides several experimentally assessed values. We gathered records for the same protein-ligand pair into groups and calculated the confidence interval of single experimental measurement for each pair, and got the average value *CI_exp_*=1.3 order of magnitude. Also we calculated the spread across maximal and minimal values for each pair, and obtained the average value of 0.9 order (Fig B in S1 Text). These values look huge, so we performed an additional investigation on the experimental affinity measurements. We found out that existing measurement techniques actually have low accuracy [49],and the obtained values strongly depend on the experiment design and human factor [49, 50]. Also in almost all cases authors do not state the binding type [51]. This means that the real precision of experiment is lower than authors often report.

Since the raw database has a number of limitations for its use as a training dataset, we applied a complex procedure of filtering, aggregation and ranking of original data to provide high-quality dataset with a maximum possible number of training instances. Also, we applied a complex approach to divide the test part of the prepared dataset including cold target and drug scaffold splits and made the resulting dataset freely available (see Data Availability section).

We trained a neural network model with the prepared BindingDB dataset and got metrics values RMSE(pIC50)=0.66 (Fig 5D) and RMSE(pKi)=0.67 (Fig 6E). Also, here we report the predicted values on the test dataset in comparison with target values for both pKi and pIC50 measurements (see Data Availability section). From our results follows that the model tends to overestimate low affinities and underestimate high ones and the most affected are the low affinity cases (pKi*<*5). However, we consider this issue non-relevant because practically such affinity ranges are less important for researchers. The average confidence interval of the model prediction on a large dataset can be estimated as *CI_calc_* = 1.96 *·RMSE* = 1.3 (see Materials and Methods). This means that our model yields the confidence level comparable with a single experimental measurement (Fig 5F). Thus, our model reached a confidence level comparable to training dataset, supporting our proposition that the primary way to tackle computational prediction of the drug-target affinity relies mainly on high quality data, and neural network architecture is less important.

In order to additionally validate the DrugForm-DTA model predictions, we performed a molecular modeling test for 127 protein-ligand pairs selected from the test set. We applied three different approaches based on molecular docking, molecular mechanics (MM-GBSA), and semiempirical quantum chemical method (PM7+COSMO). Also we introduced a modeling score function to compose the binding scores obtained from all methods. Molecular modeling scores demonstrate significantly lower correlation with experimental measurements (*C_Mod_*=0.20) compared to DrugForm-DTA model affinity predictions (*C_DT A_*=0.76). High resolution (*<*1.5Å) and amino acid sequence completeness are the keystones of molecular modeling, and none of our selected proteins have them. Moreover, molecular docking usually yields qualitative but not quantitative binding affinity estimation, and equally high modeling score values may correspond to ligands with a wide range (from 5 to 8) of experimental pKi/pIC50 values.

DrugForm-DTA model does not require 3D-structure of the protein, but uses only the primary amino acid sequence. However, molecular modeling methods can be useful for predicting spatial binding of a ligand to a protein, which complements rather than replaces the use of the DTA model.

We state that high-quality data is a primary source to train an accurate model. Thus, the future work on improving the DTA model quality is enhancing the training dataset. There are approaches to enrich the training datasets with artificial data, obtained by docking or other computational approaches [52]. Despite the fact we can get pretty huge datasets with these approaches, we consider training on real data significantly more valuable. Using large-scale artificial datasets will expectably lead to better training metrics, but we expect the behavior of the trained model to be far from reality. In this work we demonstrated that molecular modeling calculations have low correlation with experimental values. Moreover, using artificial protein structures in docking, calculated with AlphaFold2 or analogs, brings even larger deviation from real values. We performed a short test to compare docking results obtained from artificial and experimental protein structures, and found significant difference between them (see the AlphaFold2 docking comparison chapter in S1 Text). This also means that our model is expected to perform better on the proteins, existing in the BindingDB. Despite ESM’s ability to process any proteins, the DTA model is trained on a certain list of proteins.

To sum it up, we created a filtered and refined drug-target affinity dataset from the BindingDB database. We selected a modestly simple but yet effective neural network structure and validated it on standard benchmarks. After that we trained a high performance DTA model with this neural network at the refined BindingDB dataset. We also performed an additional validation of the obtained DTA model with molecular modeling methods.

## Materials and methods

### DTA model training

#### SMILES

The SMILES (Simplified Molecular Line Entry System) notation is a string representation of a molecular graph [7], both machine- and human-readable. SMILES has become one of the standards for representing the molecular structure in chemoinformatics. On the one hand, SMILES is easy to convert into molecular structure, unlike the human-readable IUPAC (International Union of Pure and Applied Chemistry) name [53]. The possibility of converting SMILES into a traditional IUPAC name has also been shown [54]. On the other hand, unlike the machine-readable InChI (International Chemical Identifier) notation [55], SMILES explicitly represents a molecular graph, and small changes in the structure lead to small changes in the SMILES notation, which makes SMILES an optimal option for representing a molecule in machine learning approaches.

#### Training-test split

It is common to separate a test part (subset) from the dataset, which does not participate in training of the ML model directly or indirectly, but is used to assess the quality of the model after training. The standard way to divide a dataset into training and testing subsets is random splitting. In the DTA task, there is a possibility that individual proteins or ligands are close to each other, but nevertheless get into different parts of the split. This can cause an effect similar to overfitting - the quality of the resulting model will be overestimated. Researchers traditionally use the following methods to strengthen the requirements for separating the test subset:

- Cold target split - test examples are selected with proteins that were not in the training set;
- Cold drug split - test examples are selected with ligands that were not in the training set;
- Drug scaffold split - test ligands are selected that have the rarest structural frameworks - scaffolds.

Cold drug split is a simpler to implement option for dividing ligands into groups, but does not take into account the structural proximity of molecules [56]. Drug scaffold split is more complex. A scaffold is built for each molecule, after which the examples are sorted by the frequency of the scaffold. Records with the rarest scaffolds represent the most unique molecules and are included in the test set.

In this work, we used a combination of approaches: forming half of the test set using the cold target split, the other half - using the drug scaffold split. Thus, a balanced test dataset was obtained, which represents a fairly complex challenge for the model, sensitivity to ligands and proteins differences.

#### Transformer

Transformer-like neural networks are widely used in various ML tasks, allowing to achieve outstanding results in different areas of computer science: natural language processing (NLP), computer vision (CV), etc. The original Transformer is presented by Vaswani et al. [32] as a tool for solving the machine translation task (Seq2Seq). The main idea is to overcome the main problem of recurrent networks - the vanishing gradient problem. Thanks to the trained attention mechanism, the model can possess complete information about the entire data sequence at any moment. Self-attention identifies which elements of the input sequence are relevant in relation to its other elements. Decoder-encoder attention identifies which elements of the input sequence are relevant for inference of the next element in the output sequence.

The original Transformer was used to develop the BERT (Bidirectional Encoder Representations from Transformers) [57] and GPT (Generative Pre-trained Transformer) [58] architectures, various variations of which are used by researchers in most modern machine learning tasks.

#### Chemformer

We use Chemformer to encode ligand molecules into embeddings [33]. It is a transformer-like model based on BART (Bidirectional Auto-Regressive Transformer) [59], a combination of BERT and a transformer-decoder. The Chemformer model is pre-trained on a large set (100 millions) of molecules represented as SMILES. The “combined” variant was used, trained with masking in combination with SMILES augmentation. The embedding dimension size is 512.

#### ESM

To encode proteins into embeddings, we use the ESM model (variant ESM2 T30 150M UR50D) [30]. This 30-layer transformer-like model was trained on *∼*27M amino acid sequences and is primarily designed to predict 3D protein structures from a given sequence. The language model underlying ESM, unlike another well-known protein structure prediction model, AlphaFold [29], has comparable accuracy but requires less computation time. In addition, AlphaFold2 and other alternative models use the multiple sequence alignment (MSA) to achieve optimal performance. However, ESM generates structure predictions using internal representations of the language model and requires only the primary amino acid sequence as input. The embedding size is 640.

#### Other tools

All program code is written in Python using the PyTorch framework version 1.13 . The implementation of transformer layers is taken from the fairseq library [60]. We also used RDKit v2023.9.5 and MolVS v0.1.1 to process and standardize SMILES. NVidia A100 GPU was used for training. We used a multitask training procedure with missed values. Both values for one protein-ligand complex are not always available in the dataset, so the training loss function was modified in order to not influence those weights which are connected to the missing output target. Since the pKi and pIC50 values correlate, training one of the parameters improves the prediction of the second parameter.

#### Benchmark metrics

MSE (Mean Squared Error) is a basic regression metric, used in benchmarks. RMSE (Root Mean Squared Error) is calculated as a square root of MSE. Both metrics could be recalculated into each other. Still, RMSE has a remarkable feature to reflect the distance from the target values, because they have the same scale. Thus we used RMSE to estimate performance of the trained model.

C-Index (Concordance Index) is a measure of the rank correlation between the predicted risk values and the observed time points [61]. Originally it is a classification metric, but it could be adapted to regression tasks. To calculate the metric values we used the code from the Affinity2Vec work [62]. We followed the established tradition in DTA works, but we see this metric as a questionable choice for a pure regression task.

In this work we compare our benchmark results with the numbers, given in the HGTDP-DTA [27] work, where authors collected and organized into a table the Davis and KIBA benchmark tests on a large number of other DTA models.

#### Confidence interval

This paper uses confidence interval estimations in several cases using both the Student criterion and the bootstrap algorithm. This paper uses confidence interval estimates. The first method involves coefficients from the t-distribution table: 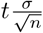. The bootstrap algorithm estimates the distribution of an estimator by resampling (often with replacement) one’s data or a model estimated from the data. Its advantage is not requiring assumptions about the data distribution. In this paper, the confidence interval of the experimental affinity constants values is calculated in both ways. In addition, through the Student’s distribution we can estimate the confidence interval of the trained affinity model forecasts. We use the RMSE metric value on the test dataset as an estimation of the standard deviation: *σ* = *RMSE*. The value of the reliability parameter of the confidence interval is the same in all cases: 95% (2*σ*), the value of the Student’s coefficient t=1.96.

### DTA benchmark datasets

To evaluate the ability of the DrugForm-DTA network to solve drug-target affinity tasks, the commonly used Davis and KIBA datasets were used.

#### Davis

The Davis dataset is a regression dataset that includes 68 drugs and 279 unique protein kinases [2]. In total, Davis contains quantitative Kd values for 25,772 ligand-protein pairs, which are calculated using the formula:

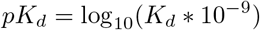

This dataset is traditionally used as the main benchmark in DTA tasks. The dataset was preliminarily divided into a training and a test set, according to the instructions of the benchmark authors. Drug molecules are presented as SMILES, protein amino acid sequences are obtained from the UniProt database [38]. The Davis dataset values distribution is presented at Fig A in S1 Text.

#### KIBA

The KIBA (Kinase Inhibitor Bioactivity) dataset is a regression dataset that includes 2,068 ligands and 229 unique protein kinases [3]. In total, KIBA contains quantitative binding values for 117,657 ligand-protein pairs. A distinctive feature of the dataset is the unification of different affinity values (Ki, IC50) into a common entity - KIBA score, which is a modified Ki:

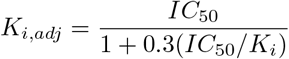

This dataset is one of the standard ones traditionally used as a DTA benchmark. The dataset was preliminarily divided into a training and a test set. The drug molecules are presented as SMILES, the protein amino acid sequences were obtained from the UniProt database [38].

### BindingDB

The DrugForm-DTA model was trained for practical application using the BindingDB database.

#### Dataset description

BindingDB is a curated dataset launched in 2000 and updated weekly [4]. It collects affinity constant values from papers, patents, and other sources, including user-uploaded ones. Among other large datasets such as PDBBind [5] or Binding MOAD [6], BindingDB is the largest, containing 2,875,634 protein-ligand pairs (1,238,443 ligands and 6,523 proteins), version “ALL 202406”.

#### Dataset preparation

The downside of the large size of the BindingDB dataset is the significant amount of invalid data and records not related to human organism. In addition, the dataset often has more than one measurement available for a single protein-ligand complex, making it impossible to train a model without prior data adaptation. We used a complex filtering procedure to prepare the dataset for training. We inserted missing UniProt IDs and obtained full amino acid sequences, then applied mutations, given in the BindingDB record, to the sequences obtained from UniProt [38]. We filtered the records: without affinity values, from inappropriate organisms, obtained at non-physiological temperature and acidity values, with invalid SMILES, with too large and too small proteins. We also clipped values outside the relevant range (out of 99th percentile) and normalized the values to log scale. In addition, we performed an aggregation procedure to unite values for identical protein-ligand complexes. Each pair was weighted according to the degree of confidence in the value and the value of the example. A complete representation of the filtering procedure is presented in Fig 7, indicating how many records does each operation reduce the dataset.

**Fig 7.**
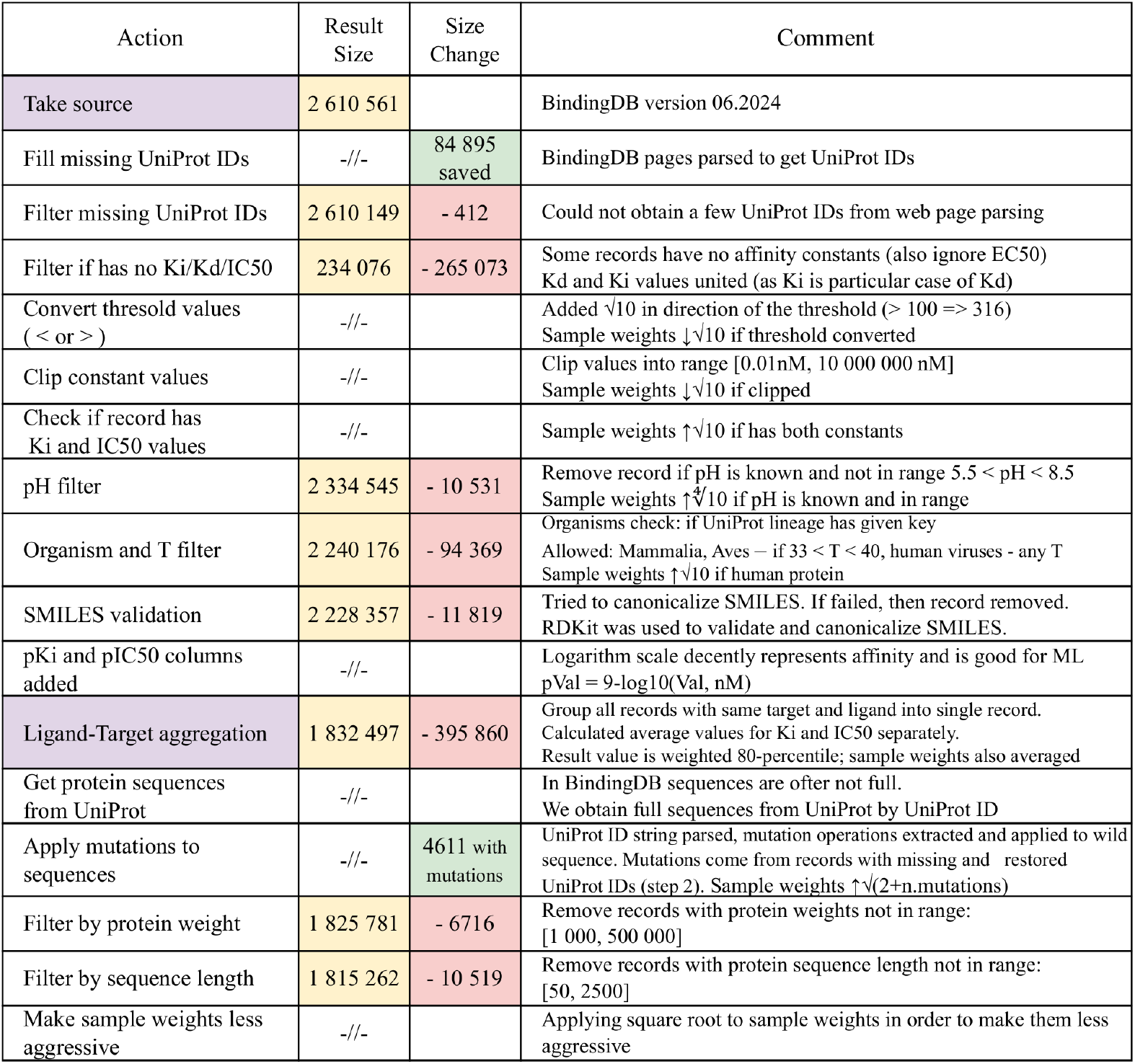
The BindingDB dataset preparation pipeline.

Here we describe some of these steps. For *∼*85,000 records from the BindingDB database, the UniProt ID of the protein is not specified, so the link to the UniProt ID was extracted from the BindingDB website [4]. The UniProt IDs were restored for 4611 unique protein-ligand complexes, including the mutant proteins.

The BindingDB dataset contains the Kd, Ki and IC50 affinity constants separately. We combined the constants Kd and Ki into a common column, but left IC50 as a separate target. We can consider Ki as a special case of Kd, but the Ki and IC50 constants, although close numerically, have different nature, so we cannot unite them into a single entity [63].

We transformed threshold values (for example, *>*5, *<*6) into exact ones based on a 10x dilution hypothesis [51, 63]. To convert the threshold value into the exact one we shift it by half of the dilution step towards the threshold side, meaning multiplying or dividing by 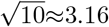. For example, the threshold value Ki *≥*10^4^ is converted to the exact value 3.16 *·* 10^4^, while simultaneously reducing the weight (importance) of this sample in training. Thus, instead of *∼* 75,000 records with pIC50=6 in the final dataset, we get *∼* 37,000 of pIC50=6, *∼* 19,000 of pIC50=5.5 and *∼* 19,000 of pIC50=6.5. For the sake of training stability and analysis simplicity, we convert them to a logarithmic scale: *pKi* = 9 −*log*_10_(*Ki, nM*), *pIC*50 = 9 −*log*_10_(*IC*50*, nM*).

If multiple records for one protein-ligand pair are available, we aggregate them into a single record with pKi and pIC50 values calculated as 80th percentile of all available measurements [37]. Three hundred randomly selected examples of visualizing 80th percentile calculation can be downloaded by the link given in the Data Availability section. These figures also contain the value spreads and the confidence intervals of the experimental measurement (95%).

Thus along with filtering and transforming the affinity values, we obtained their weights - importance for the model training. Initially, all weights were equal to “1”. During dataset processing, the record weight was multiplied by a certain factor if it is important, or divided by it if the significance of the record is questionable. A typical value of the factor for a significant criterion is 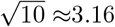, i.e. 1/2 order, and 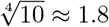 (1/4 order) for an insignificant criterion. A special case is increasing the importance of records with mutations. The more mutations are in the protein, the more valuable this example is. Therefore, the increasing coefficient for them is 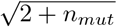, where *n_mut_* is the number of mutations in the protein. We analyzed the resulting distribution of weights and decided that it is too strong (the range of values is from 0.1 to 10), so we lightened it out using the square root operation and got the range of weights from 0.3 to 3.

#### BindingDB selective subset

To conduct detailed testing of the DTA model, we manually selected a small subset of protein-ligand complexes from samples not involved in model training. This set contains 7 proteins: melatonin receptor 1A (MTR1A), melatonin receptor 1B (MTR1B), orexin/hypocretin receptors type 1 (OX1R) and type 2 (OX2R), blood coagulation factor XIa (FA11), epidermal growth factor receptor (EGFR), sodium- and chloride-dependent glycine transporter 1 (SC6A9). These proteins are targets for the creation of antitumor drugs, drugs for insomnia and depression, as well as anticoagulants. We chose the ligands for them from the BindingDB test set (scaffold split) with affinity values in a wide range for positive and negative control. As a result, 7 ligands were selected for the MTR1B protein, and 20 ligands for the other proteins. Thus, the obtained selective subset includes 127 protein-ligand complexes.

### Molecular modeling methods

We performed validation of the affinity model using molecular modeling methods on a selective subset of 127 protein-ligand complexes. Virtual screening of this subset included molecular docking, calculation of binding energy using molecular mechanics and semiempirical quantum chemistry evaluation.

#### Molecular docking

We have chosen crystal structures of the proteins in the selective subset from the RCSB Protein Data Bank (PDB ID: 4ZJ8, 6TPN, 8A27, 4CRC, 6ME2, 6ZBV, 6ME6) [44]. All structures were obtained by X-ray diffraction, structures with good resolution (*<*2Å) and no missing atoms and amino acid residues were a priority. Molecular docking and pre-preparation of proteins and ligands were performed using Schrödinger Suite [64]. Protein structures were prepared using Protein Preparation Wizard [65], and ligand structures were prepared using the LigPrep tool [66] taking into account stereoisomers and ring conformations. Proteins and ligands were parameterized in the OPLS force field [67]. Molecular docking was performed using the Glide docking program [45]. A potential grid with the size of 20×20×20Å was pre-built for each receptor. Glide performs a complete systematic search of the conformational, orientational and positional space of the ligand in the protein active site. In this search, an initial coarse positioning and scoring phase, which dramatically narrows the search space, is followed by torsionally flexible energy optimization on an OPLS-AA non-bonded potential grid for a few hundred surviving candidate poses. The selection of the best docked ligand pose uses a model energy function that combines empirical and force field-based values. Molecular docking was performed for each ligand conformer in XP (extra precision) mode, and the ligand with the best Glide score values was used for subsequent modeling steps.

#### Molecular mechanics

We calculated the free energy of protein-ligand binding (Δ*G_bind_*) using molecular mechanics algorithms. The calculations were performed by molecular mechanics methods using the generalized Born model MM-GBSA (General Born Surface Area) [46] implemented in the Prime Schrödinger software package [68]. To calculate the binding energy of MM-GBSA, the position of the ligand in the protein active site obtained as a result of Glide docking was used. The binding energy Δ*G_bind_* was calculated using the formula:

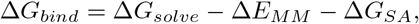

where Δ*G_solve_* — is the difference in the solvation energy of the GBSA protein-ligand complex and the sum of the free protein and free ligand solvation energies, Δ*E_MM_* - is the difference in minimized energies between the protein-ligand complex and the sum of the free protein and free ligand energies, Δ*G_SA_* - is the difference in the surface area energies of the complex and the sum of the free protein and free ligand surface area energies. The method also allows the calculation of the ligand deformation energy by placing the ligand in solution using the VSGB 2.0 solvation model. In addition, the MM-GBSA energy calculation allows the estimation of individual energy contributions such as electrostatic interaction energies, van der Waals interaction energies, lipophilicity and covalent interaction energies, as well as corrections for hydrogen bonds, *π* −*π* stacking, to the total binding energy.

MM-GBSA binding free energy calculations have several advantages. They are more theoretically rigorous than the empirical estimates used in molecular docking, and at the same time less computationally expensive than binding free energy modeling.

#### Quantum chemistry methods

We used semiempirical quantum chemistry methods to more accurately predict the protein-ligand binding affinity for the selective subset. Quantum chemistry methods allow eliminating false positive results that occur when using molecular docking. The protein-ligand binding enthalpy was calculated by the PM7 method [47] using the ligand position in the active site of the protein obtained as a result of molecular docking. In the first step, the energy of the protein-ligand complex was locally optimized using the PM7 method. During optimization, the coordinates of all ligand atoms were varied, and all protein atoms were fixed. Then, the energy of the complex was recalculated using PM7 and the COSMO solvent model [48] for all fixed atoms (1SCF+COSMO). The energy of unbound ligand formation is calculated similarly - local optimization of the energy of macrocyclic and non-aromatic ring conformations generated by LigPrep [66] and subsequent optimization taking into account the solvent (1SCF+COSMO) were performed. Among all ligand conformations, the enthalpy with the minimum value was chosen as the enthalpy of unbound ligand formation. The energy of the unbound protein was calculated by the 1SCF+COSMO method for a fixed protein conformation, which was used for molecular docking.

The enthalpy of binding was calculated using the equation:

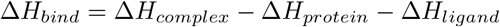

where Δ*H_complex_* is the enthalpy of optimized complex formation, Δ*H_protein_* is the enthalpy of unbound protein formation, Δ*H_ligand_* is the enthalpy of unbound ligand formation. The calculated enthalpy values were used to rank the affinity of the ligand to the protein.

#### Modeling score

We developed an algorithm to combine the results of ligand-protein binding measurements for the three approaches: molecular docking, MM-GBSA analysis, quantum chemistry methods (PM7+COSMO). This algorithm is necessary because each of the molecular modeling methods evaluates protein-ligand binding in different quantities: docking score, MM-GBSA binding energy (Δ*G_bind_*) and quantum chemistry binding enthalpy (Δ*H_bind_*).

The final ligand binding score was calculated by combining the scores of molecular docking, MM-GBSA, and the quantum chemical approach. To calculate the modeling score, a preliminary ranking procedure was performed for the interaction metrics obtained in each of the molecular modeling methods. As a result, ranges with low, medium, and high modeling scores were selected for the metrics of each method, as well as a failed category for protein-ligand pairs where modeling failed at one or more molecular modeling approaches. The final modeling score was the arithmetic mean of these scores.

## Conclusion

We developed the DrugForm-DTA model to predict the binding affinity of protein-ligand complexes. Our results indicate very high accuracy on the KIBA and Davis benchmarks (top1/top2). The DrugForm-DTA model, trained on the refined BindingDB dataset, yields the confidence level comparable with a single experimental measurement. The trained DTA model along with the benchmark models are freely available. Also, we publish the code that can run and retrain these models. The obtained BindingDB dataset with the train-test split and the building script is also available, so everyone can use it to train own models and even make a benchmark from it. We compared DrugForm-DTA with molecular modeling approaches and can conclude that accuracy of the trained neural network outperforms classical ways to assess binding of the small ligands and proteins. Our molecular modeling results are also published.

In the end, the DrugForm-DTA model is applicable for a wide range of the problems and represents a high potential method for discovery of the new therapeutic molecules for a wide range of diseases.

## Data Availability

The data and the code that support the findings of this study are openly available in Zenodo at https://doi.org/10.5281/zenodo.14949569, reference number 14949569, and in Github at https://github.com/drugform/uniqsar.

The code repository contains the framework to train and benchmark models, and also to launch them for inference. Check the readme file to get detailed usage info and instructions to reproduce the steps of this work.

The repository cannot serve big files, so they are placed at Zenodo. Follow instructions at the code repository to merge the data into your cloned repository.

Among other files, there are:

- The files data/bindingdb/bindingdb.csv and data/bindingdb/bindingdb test.csv are the cleaned BindingDB dataset we used to train and evaluate DrugForm-DTA.
- The DrugForm-DTA model itself is stored at the models/bindingdb directory.
- The selective test results.csv file is the selective test results for 127 protein-ligand pairs, containing target values from BindingDB, model predictions, and overall modeling score results.
- The file data/bindingdb/pair stats.tar.gz contains histograms of the distribution of affinity constant values for 296 individual protein-ligand pairs from BindingDB with 80th percentile values and confidence intervals plotted.
- The files models/bindingdb/test metrics/pKi.csv, models/bindingdb/test metrics/pIC50.csv contain DTA model predictions compared to targets for the test dataset.

## Supporting information

**S1 Text. Supporting information. Table A.** Glide docking results for three factor Xa inhibitor-based. **Fig A. Davis dataset.** A: pKd affinity values distribution in the Davis dataset. Almost 3/4 of all examples in the dataset have a single value of 5. Examples with other values are barely distinguishable in the histogram. B: pKd affinity values distribution in the Davis dataset when excluding pKd=5. Examples with other values become visible, but the dataset size is reduced by three times. **Fig B. Value distribution for individual protein-ligand pairs in the original BindingDB dataset.** The average spread is 0.9 orders of magnitude. There are examples with a spread of as much as 8 orders of magnitude. A significant number of pairs have zero spread, i.e. all measurements for a given pair have the same values. It is statistically unlikely that two independent experiments will give exactly the same value. These situations are presumably due to either duplication of the same result or rough rounding. If these records are excluded from the spread estimate, the average spread will increase. **Fig C. Molecular modeling EGFR.** A: 2D structures of 20 ligands for EGFR used for the selective test. Molecules with low modeling scores are marked red, and grey means modeling failed. B: Scatter plot visualizes the correlations among experimental values, DrugForm-DTA predicted values and modeling scores. All modeling failed protein-ligand complexes marked grey and related to EGFR protein (C_Mod_(EGFR)=0.18, C_DTA_(EGFR)=0.80) **Fig D. Molecular modeling MTR1A.** A: 2D structures of 20 ligands for MTR1A used for the selective test. Molecules with low modeling scores are marked red. B: Scatter plot visualizes the correlations among experimental values, DrugForm-DTA predicted values and modeling scores. Modeling of MTR1A binding revealed high modeling score values for almost all ligands from the test set, even for those with lower experimental values, that leads to low correlation values (C_Mod_(MTR1A)=-0.03, C_DTA_(MTR1A)=0.80). Thus, molecular scores are poorly suitable to distinguish between high and medium affinities (pKi/pIC50 in range from 5 to 8). **Fig E. Molecular modeling OX2R.** A: 2D structures of 20 ligands for OX2R used for the selective test. Molecules with low modeling scores are marked red. B: Scatter plot visualizes the correlations among experimental values, DrugForm-DTA predicted values and modeling scores. Modeling of OX2R binding revealed high modeling score values for almost all ligands from the test set, even for those with lower experimental values, that leads to low correlation values (C_Mod_(OX2R)=-0.26, C_DTA_(OX2R)=0.70). Thus, molecular scores are poorly suitable to distinguish between high and medium affinities (pKi/pIC50 in range from 5 to 8). **Fig F. Molecular modeling OX1R.** A: 2D structures of 20 ligands for OX1R used for the selective test. Molecules with low modeling scores are marked red. B: Scatter plot visualizes the correlations among experimental values, DrugForm-DTA predicted values and modeling scores. Modeling of ligands binding with OX1R (C_Mod_(OX1R)=-0.04, C_DTA_(OX1R)=0.84) shows no correlation with experiment. Fig G. Molecular modeling FA11. A: 2D structures of 20 ligands for FA11 used for the selective test. Molecules with low modeling scores are marked red. B: Scatter plot visualizes the correlations among experimental values, DrugForm-DTA predicted values and modeling scores. For FA11 (C_Mod_(FA11)=0.06, C_DTA_(FA11)=0.55) the modeling scores do not fully correlate with experiment, that is supposedly caused by missing a significant part of the amino acid residues of the light chain. Fig H. Molecular modeling SC6A9. A: 2D structures of 20 ligands for SC6A9 used for the selective test. Molecules with low modeling scores are marked red. B: Scatter plot visualizes the correlations among experimental values, DrugForm-DTA predicted values and modeling scores. A large number of ligands with low modeling scores are observed for protein–ligand complexes with SC6A9 (C_Mod_(SC6A9)=0.27, C_DTA_(SC6A9)=0.70). **Fig I. Molecular modeling MTR1B.** A: 2D structures of 7 ligands for MTR1B used for the selective test. Molecules with low modeling scores are marked red. B: Scatter plot visualizes the correlations among experimental values, DrugForm-DTA predicted values and modeling scores. A large number of ligands with low modeling scores are observed for protein–ligand complexes with MTR1B (C_Mod_(MTR1B)=0.40, C_DTA_(MTR1B)=0.54).

## Acknowledgments

The study was funded by the Centre for Strategic Planning and Management of Biomedical Health Risks of the Federal Medical Biological Agency.

## References

1. Zhang S, Jiang M, Wang S, Wang X, Wei Z, Li Z. SAG-DTA: Prediction of Drug–Target Affinity Using Self-Attention Graph Network. International Journal of Molecular Sciences. 2021;22(16). doi:10.3390/ijms22168993.

2. Davis MI, Hunt JP, Herrgard S, Ciceri P, Wodicka LM, Pallares G, et al. Comprehensive analysis of kinase inhibitor selectivity. Nature Biotechnology. 2011;29(11):1046–1051. doi:10.1038/nbt.1990.

3. Tang J, Szwajda A, Shakyawar S, Xu T, Hintsanen P, Wennerberg K, et al. Making Sense of Large-Scale Kinase Inhibitor Bioactivity Data Sets: A Comparative and Integrative Analysis. Journal of Chemical Information and Modeling. 2014;54(3):735–743. doi:10.1021/ci400709d.

4. Gilson M, Liu T, Baitaluk M, Nicola G, Hwang L, Chong J. BindingDB in 2015: A public database for medicinal chemistry, computational chemistry and systems pharmacology. Nucleic acids research. 2015;44. doi:10.1093/nar/gkv1072.

5. Wang R, Fang X, Lu Y, Yang CY, Wang S. The PDBbind Database: Methodologies and Updates. Journal of Medicinal Chemistry. 2005;48(12):4111–4119. doi:10.1021/jm048957q.

6. Benson ML, Smith RD, Khazanov NA, Dimcheff B, Beaver J, Dresslar P, et al. Binding MOAD, a high-quality protein–ligand database. Nucleic Acids Research. 2007;36(suppl 1):D674–D678. doi:10.1093/nar/gkm911.

7. Weininger D. SMILES, a chemical language and information system. 1. Introduction to methodology and encoding rules. Journal of Chemical Information and Computer Sciences. 1988;28(1):31–36. doi:10.1021/ci00057a005.

8. Rogers D, Hahn M. Extended-Connectivity Fingerprints. Journal of Chemical Information and Modeling. 2010;50(5):742–754. doi:10.1021/ci100050t.

9. Morgan HL. The Generation of a Unique Machine Description for Chemical Structures-A Technique Developed at Chemical Abstracts Service. Journal of Chemical Documentation. 1965;5(2):107–113. doi:10.1021/c160017a018.

10. Matsuzaka Y, Uesawa Y. Optimization of a Deep-Learning Method Based on the Classification of Images Generated by Parameterized Deep Snap a Novel Molecular-Image-Input Technique for Quantitative Structure–Activity Relationship (QSAR) Analysis. Frontiers in Bioengineering and Biotechnology. 2019;7. doi:10.3389/fbioe.2019.00065.

11. Khokhlov I, Krasnov L, Fedorov MV, Sosnin S. Image2SMILES: Transformer-Based Molecular Optical Recognition Engine. Chemistry–Methods. 2022;2(1):e202100069. doi:10.1002/cmtd.202100069.

12. Zhu X, Liu J, Zhang J, Yang Z, Yang F, Zhang X. FingerDTA: A Fingerprint-Embedding Framework for Drug-Target Binding Affinity Prediction. Big Data Mining and Analytics. 2023;6(1):1–10. doi:10.26599/BDMA.2022.9020005.

13. Jiang M, Li Z, Zhang S, Wang S, Wang X, Yuan Q, et al. Drug–target affinity prediction using graph neural network and contact maps. RSC Adv. 2020;10(35):20701–20712. doi:10.1039/D0RA02297G.

14. Yang Z, Zhong W, Zhao L, Yu-Chian Chen C. MGraphDTA: deep multiscale graph neural network for explainable drug–target binding affinity prediction. Chem Sci. 2022;13(3):816–833. doi:10.1039/D1SC05180F.

15. Qian Y, Ni W, Xianyu X, Tao L, Wang Q. DoubleSG-DTA: Deep Learning for Drug Discovery: Case Study on the Non-Small Cell Lung Cancer with EGFRT790M Mutation. Pharmaceutics. 2023;15(2). doi:10.3390/pharmaceutics15020675.

16. Nguyen T, Le H, Quinn TP, Nguyen T, Le TD, Venkatesh S. GraphDTA: predicting drug–target binding affinity with graph neural networks. Bioinformatics. 2020;37(8):1140–1147. doi:10.1093/bioinformatics/btaa921.

17. Ye H, Song Y, Wang B, Wu L, He S, Bo XC, et al. HSGCL-DTA: Hybrid-scale Graph Contrastive Learning based Drug-Target Binding Affinity Prediction. 2023; p. 947–954. doi:10.1109/ICTAI59109.2023.00142.

18. Öztürk H, Özgür A, Ozkirimli E. DeepDTA: deep drug–target binding affinity prediction. Bioinformatics. 2018;34(17):i821–i829. doi:10.1093/bioinformatics/bty593.

19. Kroll A, Ranjan S, Lercher MJ. A multimodal Transformer Network for protein-small molecule interactions enhances predictions of kinase inhibition and enzyme-substrate relationships. PLOS Computational Biology. 2024;20(5):1–23. doi:10.1371/journal.pcbi.1012100.

20. Zhang L, Wang CC, Chen X. Predicting drug–target binding affinity through molecule representation block based on multi-head attention and skip connection. Briefings in Bioinformatics. 2022;23(6):bbac468. doi:10.1093/bib/bbac468.

21. Hua Y, Song X, Feng Z, Wu X. MFR-DTA: a multi-functional and robust model for predicting drug–target binding affinity and region. Bioinformatics. 2023;39(2):btad056. doi:10.1093/bioinformatics/btad056.

22. Chen H, Li D, Liao J, Wei L, Wei L. MultiscaleDTA: A multiscale-based method with a self-attention mechanism for drug-target binding affinity prediction. Methods. 2022;207:103–109. doi:10.1016/j.ymeth.2022.09.006.

23. Chu Z, Huang F, Fu H, Quan Y, Zhou X, Liu S, et al. Hierarchical graph representation learning for the prediction of drug-target binding affinity. Information Sciences. 2022;613:507–523. doi:10.1016/j.ins.2022.09.043.

24. Voitsitskyi T, Stratiichuk R, Koleiev I, Popryho L, Ostrovsky Z, Henitsoi P, et al. 3DProtDTA: a deep learning model for drug-target affinity prediction based on residue-level protein graphs. RSC Adv. 2023;13(15):10261–10272. doi:10.1039/D3RA00281K.

25. Kalemati M, Zamani Emani M, Koohi S. BiComp-DTA: Drug-target binding affinity prediction through complementary biological-related and compression-based featurization approach. PLOS Computational Biology. 2023;19(3):1–28. doi:10.1371/journal.pcbi.1011036.

26. Ma W, Zhang S, Li Z, Jiang M, Wang S, Guo N, et al. Predicting Drug-Target Affinity by Learning Protein Knowledge From Biological Networks. IEEE Journal of Biomedical and Health Informatics. 2023;27(4):2128–2137. doi:10.1109/JBHI.2023.3240305.

27. Xiao X, Wang W, Xie J, Zhu L, Chen G, Li Z, et al. HGTDP-DTA: Hybrid Graph-Transformer with Dynamic Prompt for Drug-Target Binding Affinity Prediction. 2024;doi:10.48550/arXiv.2406.17697.

28. Mikolov T, Chen K, Corrado G, Dean J. Efficient Estimation of Word Representations in Vector Space; 2013. Available from: https://arxiv.org/abs/1301.3781.

29. Jumper J, Evans R, Pritzel A, Green T, Figurnov M, Ronneberger O, et al. Highly accurate protein structure prediction with AlphaFold. Nature. 2021;596(7873):583–589. doi:10.1038/s41586-021-03819-2.

30. Lin Z, Akin H, Rao R, Hie B, Zhu Z, Lu W, et al. Evolutionary-scale prediction of atomic-level protein structure with a language model. Science. 2023;379(6637):1123–1130. doi:10.1126/science.ade2574.

31. Ahmad W, Simon E, Chithrananda S, Grand G, Ramsundar B. ChemBERTa-2: Towards Chemical Foundation Models; 2022. Available from: https://arxiv.org/abs/2209.01712.

32. Vaswani A, Shazeer N, Parmar N, Uszkoreit J, Jones L, Gomez AN, et al. Attention Is All You Need. CoRR. 2017;abs/1706.03762.

33. Irwin R, Dimitriadis S, He J, Bjerrum EJ. Chemformer: a pre-trained transformer for computational chemistry. Machine Learning: Science and Technology. 2022;3(1):015022. doi:10.1088/2632-2153/ac3ffb.

34. Kim Y, Jeong Y, Kim J, Lee EK, Kim WJ, Choi IS. MolNet: A Chemically Intuitive Graph Neural Network for Prediction of Molecular Properties; 2022. Available from: https://arxiv.org/abs/2203.09456.

35. Bishop CM. Neural networks for pattern recognition. Oxford University Press, USA; 1995.

36. Huang K, Fu T, Gao W, Zhao Y, Roohani Y, Leskovec J, et al. Therapeutics Data Commons: Machine Learning Datasets and Tasks for Drug Discovery and Development; 2021. Available from: https://arxiv.org/abs/2102.09548.

37. Bastos M, Abian O, Johnson CM, Ferreira-da Silva F, Vega S, Jimenez-Alesanco A, et al. Isothermal titration calorimetry. Nature Reviews Methods Primers. 2023;3(1):17. doi:10.1038/s43586-023-00199-x.

38. Coudert E, Gehant S, de Castro E, Pozzato M, Baratin D, Neto T, et al. Annotation of biologically relevant ligands in UniProtKB using ChEBI. Bioinformatics. 2022;39(1):btac793. doi:10.1093/bioinformatics/btac793.

39. Wang J, Wen N, Wang C, Zhao L, Cheng L. ELECTRA-DTA: a new compound-protein binding affinity prediction model based on the contextualized sequence encoding. Journal of Cheminformatics. 2022;14. doi:10.1186/s13321-022-00591-x.

40. Hu Z, Liu W, Zhang C, Huang J, Zhang S, Yu H, et al. SAM-DTA: a sequence-agnostic model for drug–target binding affinity prediction. Briefings in Bioinformatics. 2022;24(1):bbac533. doi:10.1093/bib/bbac533.

41. Lenselink E, Dijke N, Bongers B, Papadatos G, Vlijmen H, Kowalczyk W, et al. Beyond the hype: deep neural networks outperform established methods using a ChEMBL bioactivity benchmark set. Journal of Cheminformatics. 2017;9:45. doi:10.1186/s13321-017-0232-0.

42. Feng Q, Dueva E, Cherkasov A, Ester M. PADME: A Deep Learning-based Framework for Drug-Target Interaction Prediction; 2019. Available from: https://arxiv.org/abs/1807.09741.

43. Gorriz JM, Clemente RM, Segovia F, Ramirez J, Ortiz A, Suckling J. Is K-fold cross validation the best model selection method for Machine Learning?; 2024. Available from: https://arxiv.org/abs/2401.16407.

44. Berman HM, Westbrook J, Feng Z, Gilliland G, Bhat TN, Weissig H, et al. The Protein Data Bank. Nucleic Acids Research. 2000;48:235–242. doi:10.1093/nar/28.1.235.

45. Friesner RA, Banks JL, Murphy RB, Halgren TA, Klicic JJ, Mainz DT, et al. Glide: A New Approach for Rapid, Accurate Docking and Scoring. 1. Method and Assessment of Docking Accuracy. Journal of Medicinal Chemistry. 2004;47(7):1739–1749. doi:10.1021/jm0306430.

46. Genheden S, Ryde U. The MM/PBSA and MM/GBSA methods to estimate ligand-binding affinities. Expert Opinion on Drug Discovery. 2015;10(5):449–461. doi:10.1517/17460441.2015.1032936.

47. Hostaš J, Řezáč J, Hobza P. On the performance of the semiempirical quantum mechanical PM6 and PM7 methods for noncovalent interactions. Chemical Physics Letters. 2013;568-569:161–166. doi:10.1016/j.cplett.2013.02.069.

48. Klamt A, Schüürmann G. COSMO: a new approach to dielectric screening in solvents with explicit expressions for the screening energy and its gradient. J Chem Soc, Perkin Trans 2. 1993;(5):799–805. doi:10.1039/P29930000799.

49. Jin Z, Wei Z. Molecular simulation for food protein–ligand interactions: A comprehensive review on principles, current applications, and emerging trends. Comprehensive Reviews in Food Science and Food Safety. 2024;23(1):e13280. doi:10.1111/1541-4337.13280.

50. Georgakis N, Ioannou E, Varotsou C, Premetis G, Chronopoulou EG, Labrou NE. Determination of Half-Maximal Inhibitory Concentration of an Enzyme Inhibitor. In: Labrou NE, editor. Targeting Enzymes for Pharmaceutical Development. vol. 2089. New York, NY: Springer US; 2020. p. 41–46. Available from: http://link.springer.com/10.1007/978-1-0716-0163-1_3.

51. Ren T, Zhu X, Jusko NM, Krzyzanski W, Jusko WJ. Pharmacodynamic model of slow reversible binding and its applications in pharmacokinetic/pharmacodynamic modeling: review and tutorial. Journal of Pharmacokinetics and Pharmacodynamics. 2022;49(5):493–510. doi:10.1007/s10928-022-09822-y.

52. Sadybekov AA, Sadybekov AV, Liu Y, Iliopoulos-Tsoutsouvas C, Huang XP, Pickett J, et al. Synthon-based ligand discovery in virtual libraries of over 11 billion compounds. Nature. 2022;601(7893):452–459. doi:10.1038/s41586-021-04220-9.

53. Skonieczny S. The IUPAC Rules for Naming Organic Molecules. Journal of Chemical Education. 2006;83(11):1633. doi:10.1021/ed083p1633.

54. Krasnov L, Khokhlov I, Fedorov MV, Sosnin S. Transformer-based artificial neural networks for the conversion between chemical notations. Scientific Reports. 2021;11(1):14798. doi:10.1038/s41598-021-94082-y.

55. Heller SR, McNaught A, Pletnev IV, Stein S, Tchekhovskoi DV. InChI, the IUPAC International Chemical Identifier. Journal of Cheminformatics. 2015;7.

56. Nguyen TM, Nguyen T, Tran T. Mitigating cold-start problems in drug-target affinity prediction with interaction knowledge transferring. Briefings in Bioinformatics. 2022;23(4):bbac269. doi:10.1093/bib/bbac269.

57. Devlin J, Chang MW, Lee K, Toutanova K. BERT: Pre-training of Deep Bidirectional Transformers for Language Understanding. In: North American Chapter of the Association for Computational Linguistics; 2019. Available from: https://api.semanticscholar.org/CorpusID:52967399.

58. Radford A, Narasimhan K, Salimans T, Sutskever I. Improving language understanding by generative pre-training; 2018. Available from: https://www.mikecaptain.com/resources/pdf/GPT-1.pdf.

59. Lewis M, Liu Y, Goyal N, Ghazvininejad M, Mohamed A, Levy O, et al. BART: Denoising Sequence-to-Sequence Pre-training for Natural Language Generation, Translation, and Comprehension. In: Jurafsky D, Chai J, Schluter N, Tetreault J, editors. Proceedings of the 58th Annual Meeting of the Association for Computational Linguistics. Online: Association for Computational Linguistics; 2020. p. 7871–7880. Available from: https://aclanthology.org/2020.acl-main.703.

60. Ott M, Edunov S, Baevski A, Fan A, Gross S, Ng N, et al. fairseq: A Fast, Extensible Toolkit for Sequence Modeling. CoRR. 2019;abs/1904.01038.

61. Heagerty PJ, Zheng Y. Survival Model Predictive Accuracy and ROC Curves. Biometrics. 2005;61(1):92–105. doi:10.1111/j.0006-341X.2005.030814.x.

62. Thafar MA, Alshahrani M, Albaradei S, Gojobori T, Essack M, Gao X. Affinity2Vec: drug-target binding affinity prediction through representation learning, graph mining, and machine learning. Scientific reports. 2022;12(1). doi:10.1038/s41598-022-08787-9.

63. Hoare SRJ. The Problems of Applying Classical Pharmacology Analysis to Modern In Vitro Drug Discovery Assays: Slow Binding Kinetics and High Target Concentration. SLAS Discovery. 2021;26(7):835–850. doi:10.1177/24725552211019653.

64. Schrödinger Suite: Computational Platform for Molecular Discovery & Design. Maestro, Schrödinger, LLC, New York, NY. 2024;.

65. Madhavi Sastry G, Adzhigirey M, Day T, Annabhimoju R, Sherman W. Protein and ligand preparation: parameters, protocols, and influence on virtual screening enrichments. Journal of Computer-Aided Molecular Design. 2013;27(3):221–234. doi:10.1007/s10822-013-9644-8.

66. Schrödinger LigPrep: Versatile ligand preparetion tool for structure-based workflows. LigPrep, Schrödinger, LLC, New York, NY. 2024;.

67. Kahn K, Bruice TC. Parameterization of OPLS–AA force field for the conformational analysis of macrocyclic polyketides. Journal of Computational Chemistry. 2002;23(10):977–996. doi:10.1002/jcc.10051.

68. Jacobson MP, Pincus DL, Rapp CS, Day TJF, Honig B, Shaw DE, et al. A hierarchical approach to all-atom protein loop prediction. Proteins: Structure, Function, and Bioinformatics. 2004;55(2):351–367. doi:10.1002/prot.10613.

